# *In Vitro* Antibacterial Activity of Dinuclear Thiolato-Bridged Ruthenium(II)-Arene Compounds

**DOI:** 10.1101/2023.02.21.529477

**Authors:** Quentin Bugnon, Camilo Melendez, Oksana Desiatkina, Louis Fayolles Chorus de Chaptes, Isabelle Holzer, Emilia Păunescu, Markus Hilty, Julien Furrer

## Abstract

The antibacterial activity of 22 thiolato-bridged dinuclear ruthenium(II)-arene compounds was assessed *in vitro* against *Escherichia coli, Streptococcus pneumoniae* and *Staphylococcus aureus*. None of the compounds efficiently inhibited the growth of the three *E. coli* strains tested and only compound **5** exhibited a medium activity against this bacterium (MIC (minimum inhibitory concentration) of 25 μM). However, a significant antibacterial activity was observed against *S. pneumoniae*, with MIC values ranging from 1.3 to 2.6 μM for compounds **1-3**, **5** and **6**. Similarly, compounds **2**, **5-7** and **20-22** had MIC values ranging from 2.5 to 5 μM against *S. aureus.* The tested diruthenium compounds have a bactericidal effect significantly faster than that of penicillin. Fluorescence microscopy assays performed on *S. aureus* using the BODIPY-tagged diruthenium complex **15** showed that this type of metal compound enter the bacteria and do not accumulate in the cell wall of gram-positive bacteria. Cellular internalization was further confirmed by inductively coupled plasma mass spectrometry (ICP-MS) experiments. The nature of the substituents anchored on the bridging thiols and the compounds molecular weight appear to significantly influence the antibacterial activity. Thus, if overall a decrease of the bactericidal effect with the increase of compounds’ molecular weight is observed, however the complexes bearing larger benzo-fused lactam substituents had low MIC values. This first antibacterial activity screening demonstrated that the thiolato-diruthenium compounds exhibit promising activity against *S. aureus* and *S. pneumoniae* and deserve to be considered for further studies.

## 1. Introduction

The increasing occurrence of multidrug resistant (MDR) bacteria and difficult to treat infections associated with high morbidity and mortality have become important medical issues (1–3). Yearly, the number of estimated deaths caused by antibiotic resistant infections are of 33,000 in Europe and 35,000 in the United States (4). In 2019, on a global scale 1.27 million deaths were directly attributable to drug resistance (5). Rapid and efficient solutions are required to counter potential outbreaks, with terrible life costs and economical damages (1, 6). A list of the world’s leading antibiotic-resistant bacteria with an urgent demand for new antibiotics was published by WHO (World Health Organization) (7), aiming not only to favor the surveillance and control of MDR bacteria, but also to favor research of new active compounds. Antibiotic-resistant *Enterobacteriaceae* (encompassing *Escherichia coli*), *Staphylococcus aureus* as well as *Streptococcus pneumoniae* are included in the WHO’ list (7).

To efficiently face the problem of bacterial resistance to antibiotics (intrinsic and/or acquired (8)), the improvement of already existing antibiotics should be paralleled by the develop new classes of active compounds against which bacteria are less prone to develop resistances (9).

Metal complexes reached clinical trials for the treatment of various disease and some compounds show also interesting antibacterial activity (10–12). However, the use of metal-based compounds as antibiotics is not widespread and only a limited number of compounds have undergone clinical trials (13). Examples of metal complexes active against Gram-negative and Gram-positive bacteria (*E. coli* and *S. aureus*) are provided in Table S1 (13–32). Ruthenium is one of the metals considered for the development of metal-based antibiotics being generally associated to reduced toxicity (10). Some ruthenium complexes exert good activity against Gram-positive bacteria and low activity towards Gram-negative bacteria, notable exceptions being dinuclear polypyridylruthenium(II) complexes (31, 33) and ruthenium-based carbon-monoxide-releasing molecules (34–36).

Di- and trithiolato-bridged dinuclear ruthenium(II)-arene complexes are investigated for more than a decade. This type of diruthenium complexes are generally stable (37, 38), and showed promising *in vitro* and *in vivo* anticancer activity (37, 39) and *in vitro* antiparasitic activity against *Toxoplasma gondii* (40–44) and *Trypanosoma brucei* (45). Their exact mode of action has not yet been elucidated, but a profound alteration of the structure and activity of mitochondria has been identified. (40, 41, 45).

To identify further biological applications of dinuclear thiolato-bridged ruthenium(II)-arene complexes, a library of 22 compounds was screened for potential antibacterial properties. This library included both previously reported and newly synthesized compounds with significant structural diversity: dithiolato compounds, symmetric and mixed trithiolato compounds, and conjugates containing one or more carbohydrate units, short lipid chains, benzo-fused lactams and fluorophores. The minimum inhibitory concentration (MIC) values of the compounds were measured against various strains of clinically relevant Gram-negative (*E. coli*) and Gram-positive bacteria (*S. pneumoniae* and *S. aureus).* Additional experiments to evaluate the bacteriostatic and bactericidal properties of the compounds as well as their internalization and cellular localization were also performed.

## 2. Results

### 2.1. Chemistry

Based on their structural features the compounds included in this study were organized in four families.

#### 2.1.1. Di- and trithiolato diruthenium compounds **1-8** (Family 1)

Family 1 includes eight previously known diruthenium complexes (**1-8**, Figure 1), and included the neutral dithiolato compound **4**, two symmetric trithiolato diruthenium complexes **5** and **6** with benzyl and phenyl substituents on the bridging thiols, and five mixed trithiolato diruthenium complexes **1-3**, **7** and **8** in which various structural elements were varied. Some of these compounds have previously shown high anticancer and antiparasitic activity (44, 46, 47). For example, compound **1** exhibited nanomolar range IC_50_ (half maximal inhibitory concentration) values against various parasites as *Neospora caninum* (41) and *Toxoplasma gondii* (40).

**Figure 1.**
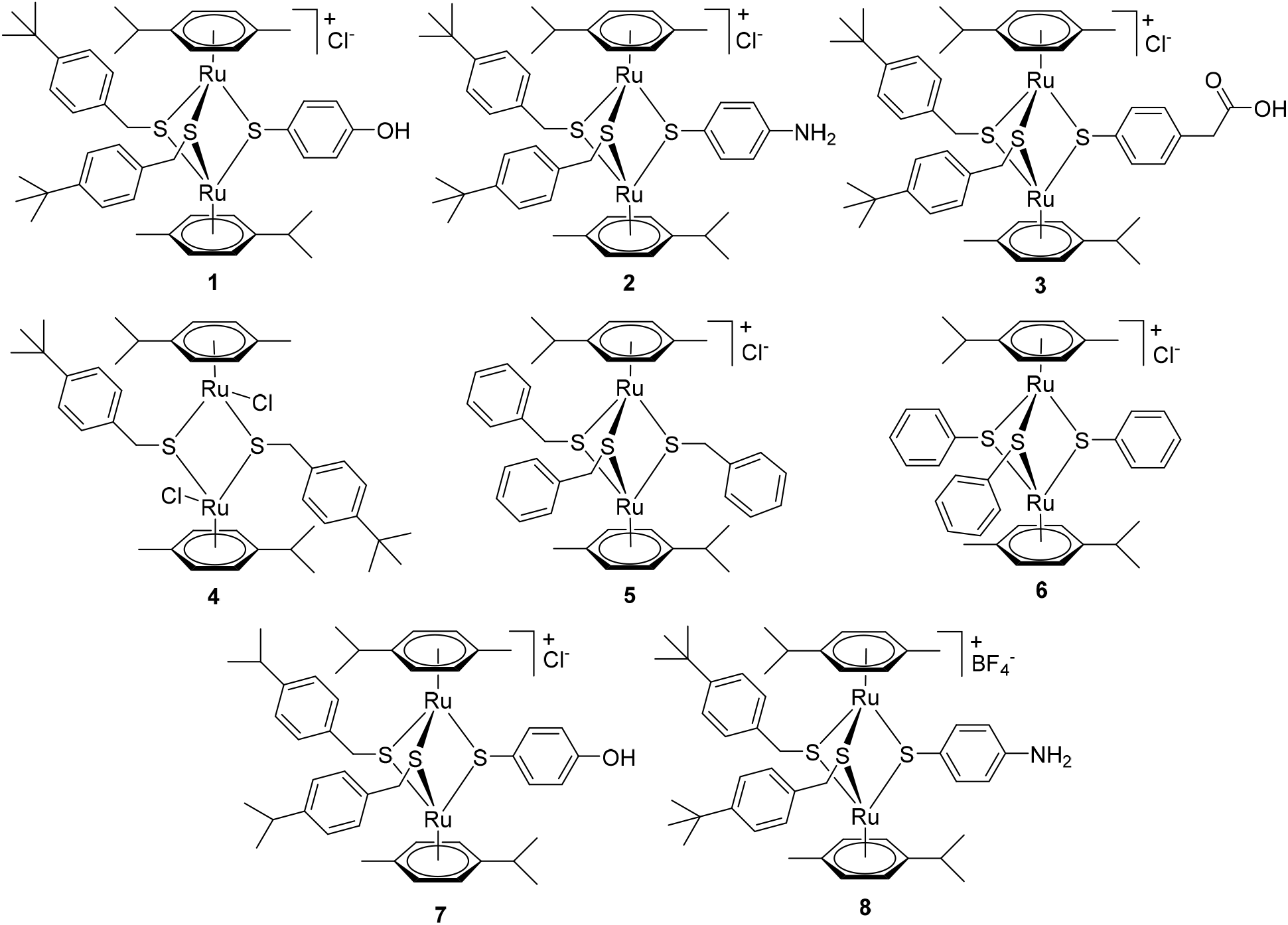
Structure of compounds **1-8** forming Family 1.

#### 2.1.2. Compounds **9-12** - diruthenium complexes conjugated to hexanoic acid and to carbohydrates (Family 2)

In this family, four trithiolato diruthenium compounds with organic molecules anchored on one of the bridge thiols were also selected. Compounds **9** and **10** (48) contain an hexanoic acid residue connected *via* an ester and, respectively, an amide bond to the diruthenium unit, and they were chosen considering that the lipophilic chain could influence membrane transfer (Figure 2).

**Figure 2.**
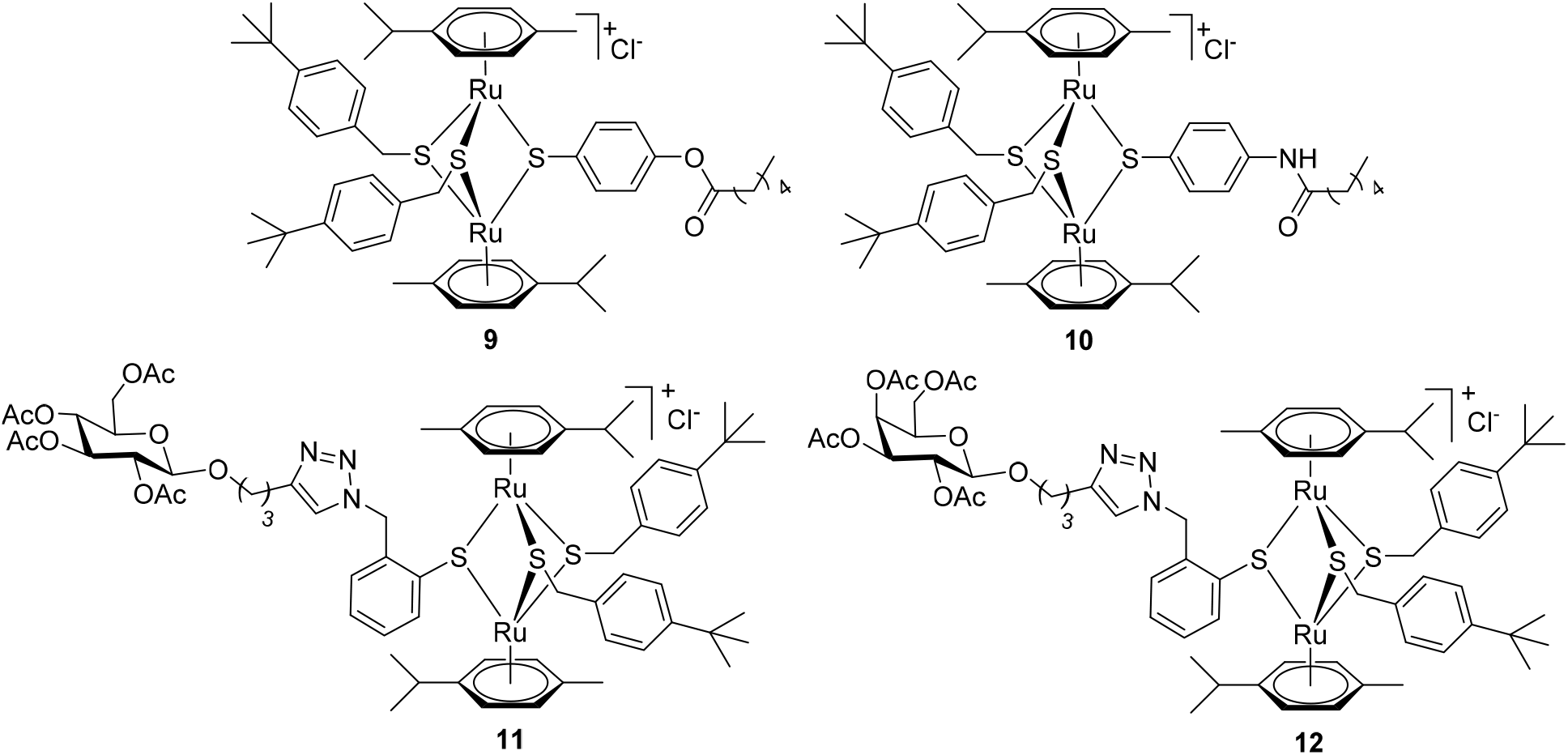
Structure of diruthenium conjugates **9-12** forming Family 2.

Compounds **11** and **12** present acetyl-protected D-glucose and, respectively D-galactose units (Figure 2), anchored to one of the bridge thiols *via* a triazole linker, and they were selected in view of a potentially facilitated bacteria internalization due to the presence of the carbohydrate unit. The synthesis, characterization, *and T. gondii* antiparasitic effects of compounds **9-12** have been previously described (49, 50).

#### 2.1.3. Compounds **13-16** - diruthenium complexes conjugated to BODIPY fluorophores (Family 3)

The development of fluorophore-labelled conjugates of metal-based drugs as traceable therapeutic agents has become extremely popular as they can provide important information relative to compounds cellular uptake, localization, and specific accumulation (42, 51). BODIPY (boron dipyrromethene, 4,4-difluoro-4-bora-3*a*,4*a*-diaza-*s*-indacene) are among the most attractive fluorophores tagged as they are generally non-toxic, photostable, and exhibit high fluorescence quantum yields (51).

Four fluorescent conjugates, **13-16**, containing a BODIPY dye unit attached to the diruthenium moiety through linkers of various lengths *via* ester (compounds **13** and **15**) or amide (compounds **14** and **16**) bonds (Figure 3) were selected. Although an important fluorescence quenching was observed after conjugating the BODIPY to the diruthenium unit, compounds **13-16** could be used as fluorescent tracers (52), and were recently investigated as potential anti-toxoplasma agents. Fluorescence microscopy investigations of these compounds showed a pattern of cytoplasmic, but not nuclear, localization in human foreskin fibroblasts (52).

**Figure 3.**
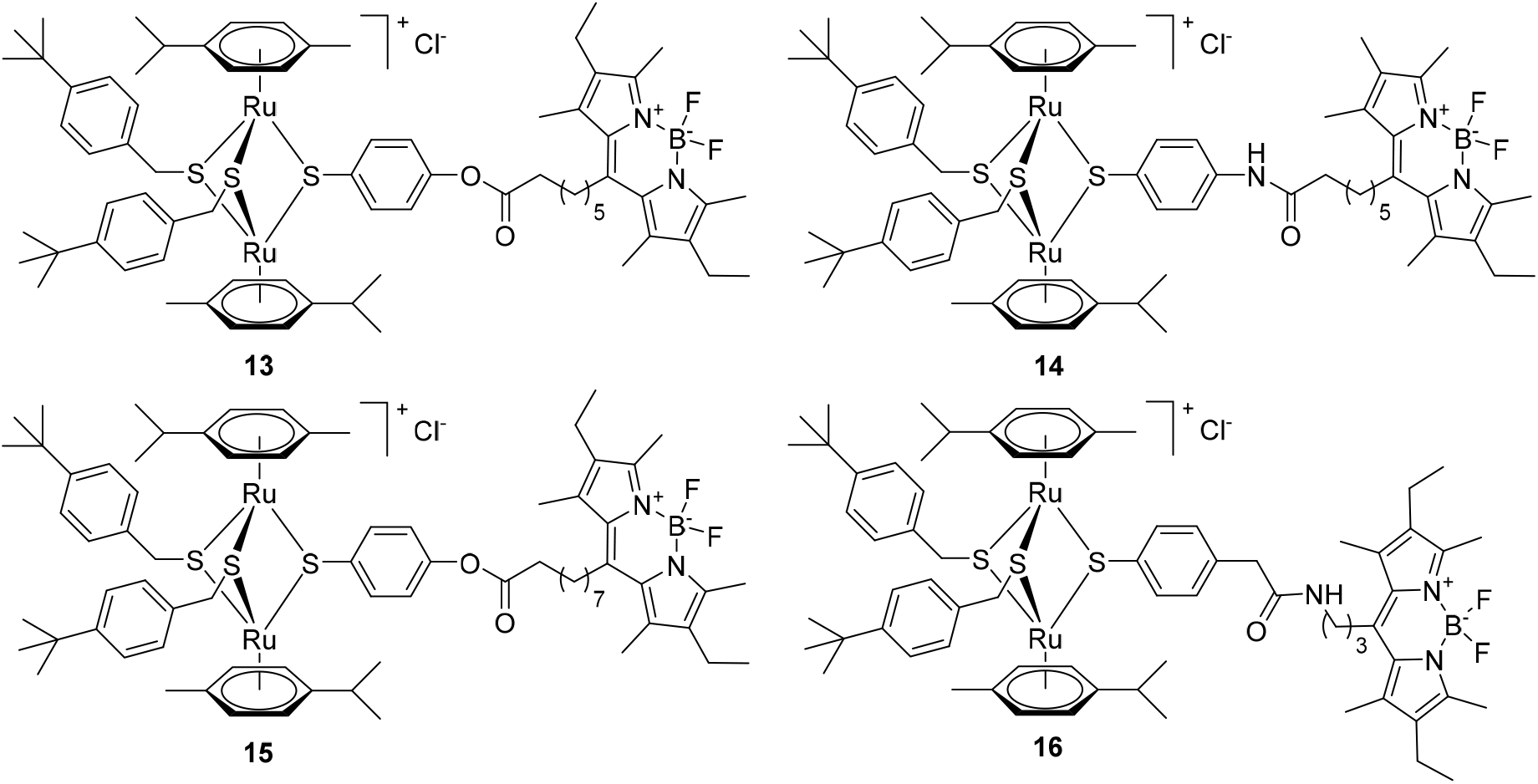
Structure of the diruthenium-BODIPY conjugates **13-16** forming Family 3.

#### 2.1.4. Compounds **17-22** – Diruthenium compounds with various thiol ligands (ortho substituted arenes and benzo-fused lactams as pending groups) (Family 4)

Six new thiolato diruthenium compounds **17-22** (Figure 4) were synthesized to evaluate structurally different ligands. The description of the synthetic protocols and the analysis and characterization of these compounds and the corresponding ligands are provided in the *Supporting information* (Schemes 1 and 2 for the synthesis of the diruthenium complexes, and Schemes S1 and S2 (*Supporting information*) for the synthesis of the thiol ligands).

**Figure 4.**
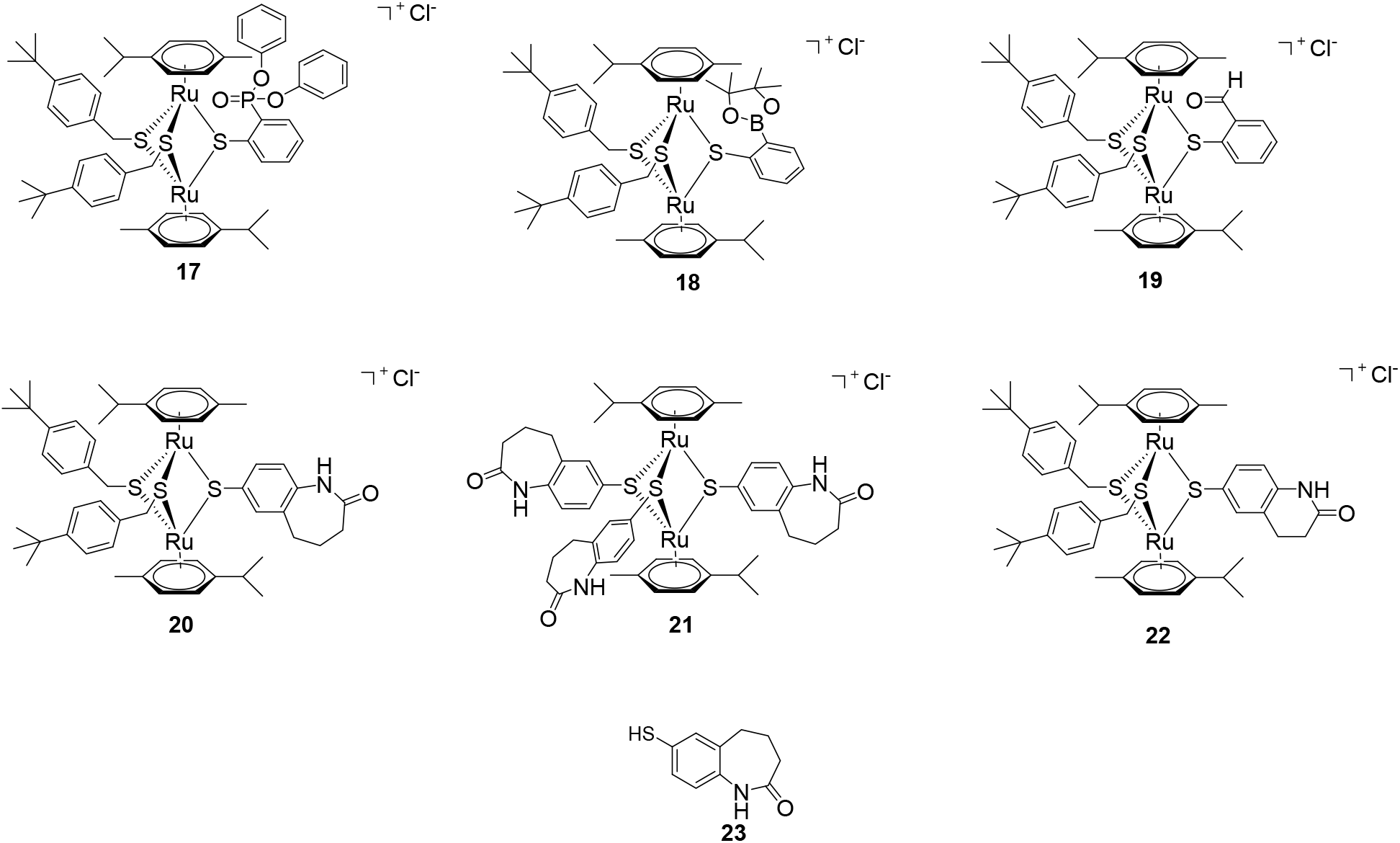
Structure of compounds **17-22** forming Family 4. **23** is the benzo-fused lactam thiol ligand used in compounds **20** and **21** and tested against the various bacteria.

In compounds **17-19** (Scheme 1), three different groups, diphenylphosphonate (**17**), 2-4,4,5,5-tetramethyl-1,3,2-dioxaborolan-2-yl (**18**), and aldehyde (**19**) were introduced in the *ortho* position rather than in *para* position of one of the bridging thiol-ligand. The synthesis of the ligands **17a** (diphenyl-(2-mercaptophenyl)phosphonate), **18a** (2-(4,4,5,5-tetramethyl-1,3,2-dioxaborolan-2-yl)benzenethiol) and **19a** (2-mercaptobenzaldehyde) (Schemes S1) are presented in detail in the Supporting information.

**Scheme 1.**
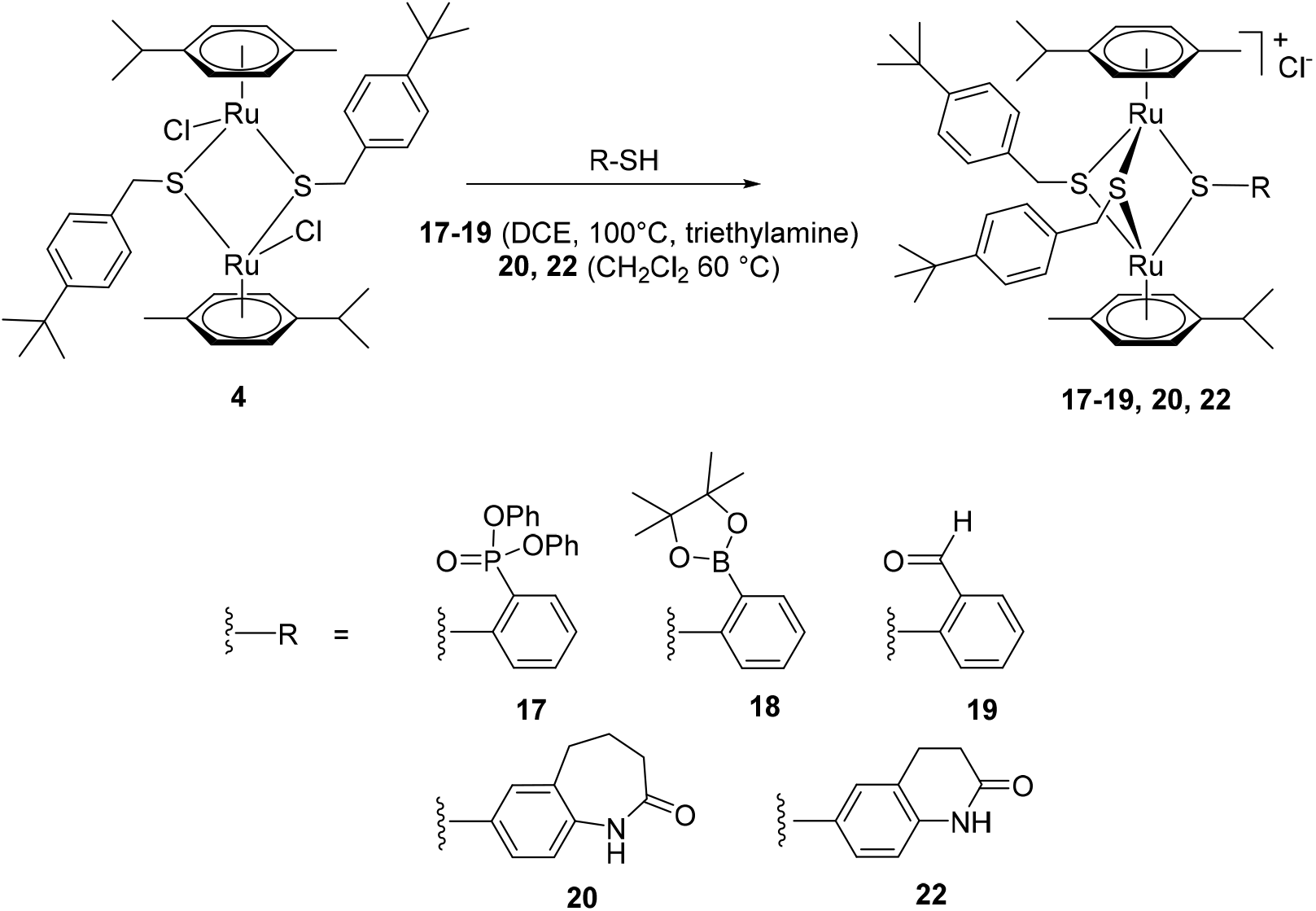
Synthesis of the mixed trithiolato diruthenium complexes **17-20** and **22**.

For compounds **20-22** (Scheme 1), two benzo-fused lactams containing thiol groups (6-mercapto-3,4-dihydroquinolin-2(1*H*)-one (**22a**) and 7-mercapto-1,3,4,5-tetrahydro-2*H*-benzo[*b*]azepin-2-one (**23**)) were used as bridging ligands instead of 4-mercaptophenol in the parent trithiolato diruthenium compound **1**. Their synthesis is provided in detail in the *Supporting information*, and were selected because benzo-fused lactams exhibit various biological activities among which antimicrobial properties (54, 55).

In the symmetric trithiolato diruthenium complex **21** (Scheme 2), the benzyl and phenyl bridging thiols in complexes **5** and **6** were replaced by three 7-mercapto-1,3,4,5-tetrahydro-2*H*-benzo[*b*]azepin-2-one units.

**Scheme 2.**
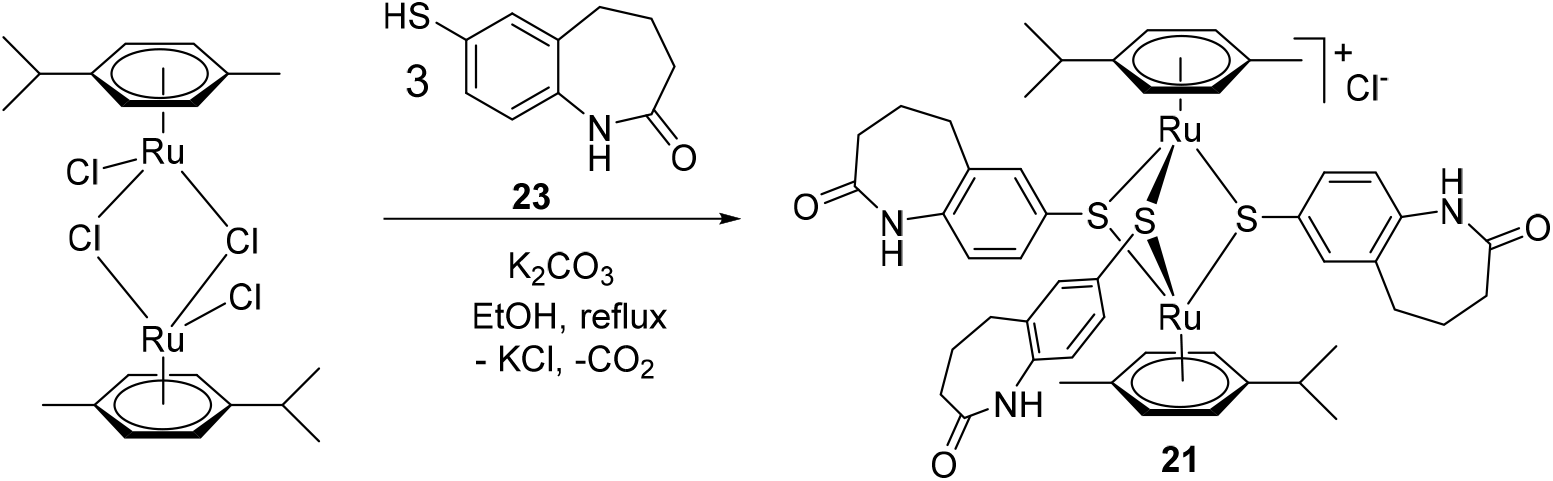
Synthesis of the symmetric trithiolato diruthenium complex **21**.

Asymmetric trithiolato diruthenium complexes **17-20** and **22** were synthetized by reacting the dithiolato intermediate **4** with an excess of the corresponding ligands **17a-19a**, **22a** and **23 (**in 1,2-dichloroethane - DCE- at 100°C or in dichloromethane at 60 °C). In the case of complexes **17-19**, excess of triethylamine was employed to ensure the completion of the reaction. All the compounds were isolated in moderate yields 41-65% (Scheme 1).

The symmetric trithiolato diruthenium complex **21** was obtained by reacting the commercially available ruthenium dimer ([Ru(*η*^6^-*p*-MeC_6_H_4_Pr^*i*^)Cl]_2_Cl_2_) with an excess of the thiol ligand **23** in refluxing EtOH in basic conditions (K_2_CO_3_) and was isolated in 56% yield (Scheme 2).

### 2.2. Stability of the diruthenium compounds

For the biological activity evaluation, 1 mM stock solutions of all compounds were prepared in dimethylsulfoxide (DMSO). ^1^H-NMR spectra of several conjugates dissolved in DMSO-*d*_6_, recorded at 25°C 5 min and 100 days after sample preparation showed no modifications, demonstrating very good stability of the diruthenium compounds in this highly complexing solvent (42, 44, 52).

Compounds **9**, **11-13** and **15** present a carboxyl ester bond that can potentially be hydrolyzed in cellular growth media. Compounds **13** and **15** as well as other similar conjugates with coumarin and BODIPY fluorescent units have been recently studied (42, 52), and for these compounds, very limited solvolysis of the ester bonds after 168 h was observed. It was concluded that diruthenium conjugates with ester linkers were stable under the conditions used for biological evaluations, and therefore, it was assumed that compounds **9**, **11-13** and **15** are sufficiently stable during the *in vitro* evaluation.

### 2.3. Determination of the MIC values on E. coli

The MIC values (lowest concentration of antibiotic at which bacterial growth is completely inhibited) of the compounds were determined on three *E. coli* strains (P53.1R (56), P54.1T (56) and P54.2R (56)), the *S. pneumoniae* D39 (NCTC 7466) (57) strain, and the *S. aureus* 20 (56) strain.

The first screening of the 22 diruthenium compounds was conducted on *E. coli* P53.1R, a colistin sensitive but extended spectrum cephalosporin resistant strain (Table 1). Most of the compounds exhibited no or very low antibacterial activity on this Gram-negative bacterium (MIC values over or at the limit of the concentration range (0.01-100 μM) used in the assays). From this library, the only compound inhibiting *E. coli* P53.1R cells growth with medium activity (MIC value of 25 μM) was the symmetric diruthenium compound **5** (Table 1). The potency of compounds **2**, **5** and **6** from Family 1 was further assessed for two additional *E. coli* strains P54.1T and P54.2R (Table 1). On the *E. coli* P54.1T and P54.2R strains, the lowest MIC values (of 25 and 20.8 μM, respectively) were obtained for the same compound **5** (Table 1), which inhibited the bacteria growth with medium efficiency compared with the reference drug colistin for which the MIC value is 0.8 μM.

**Table 1.**
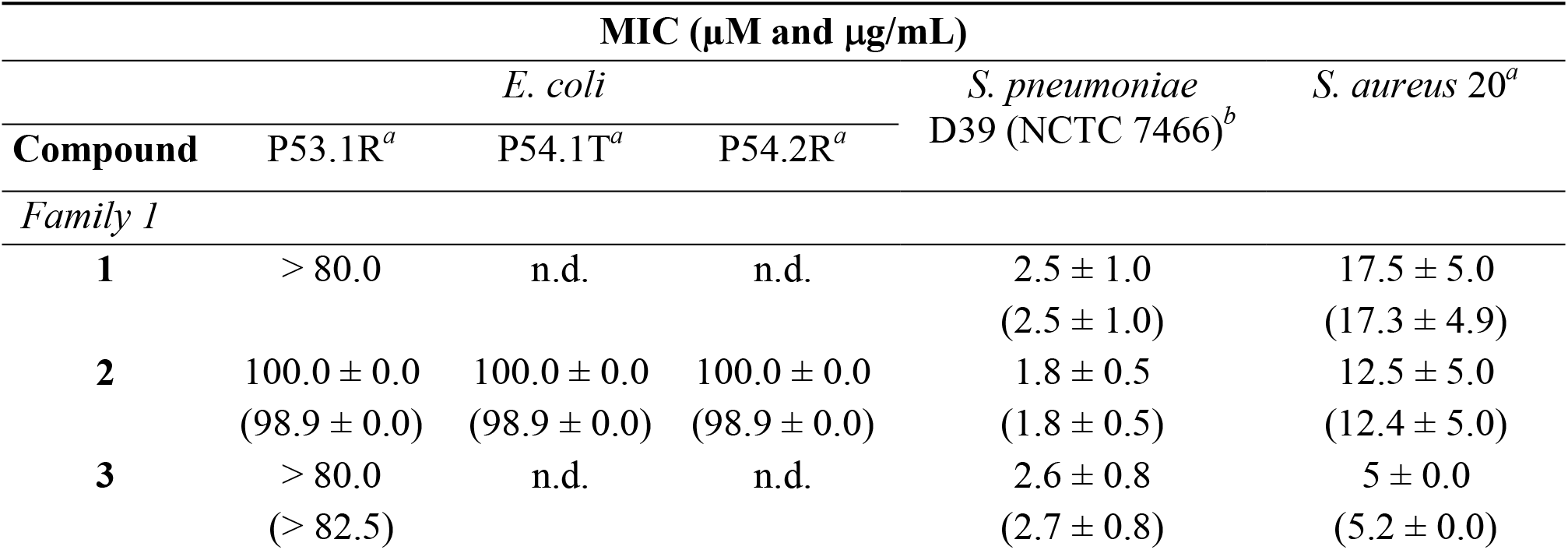
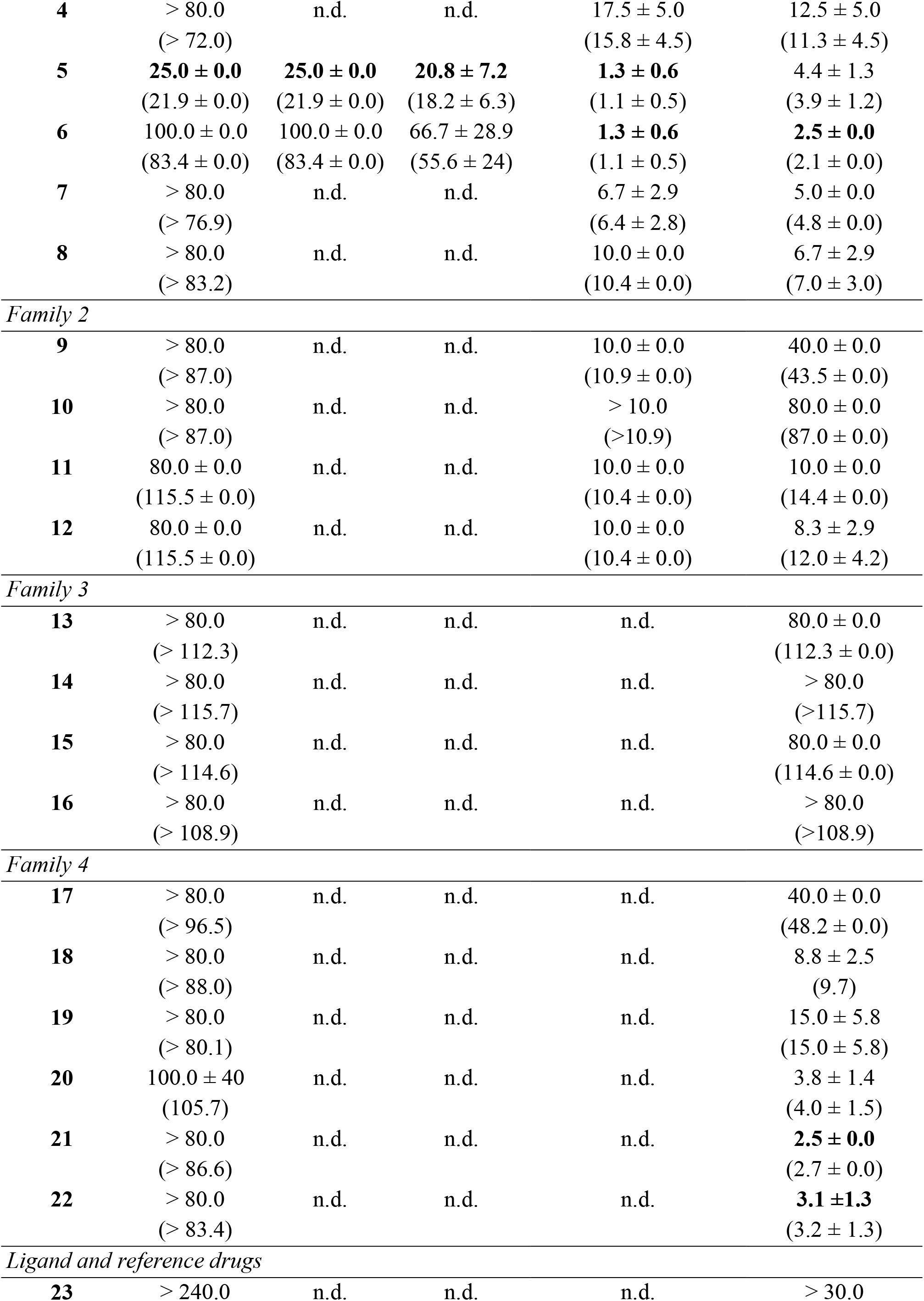
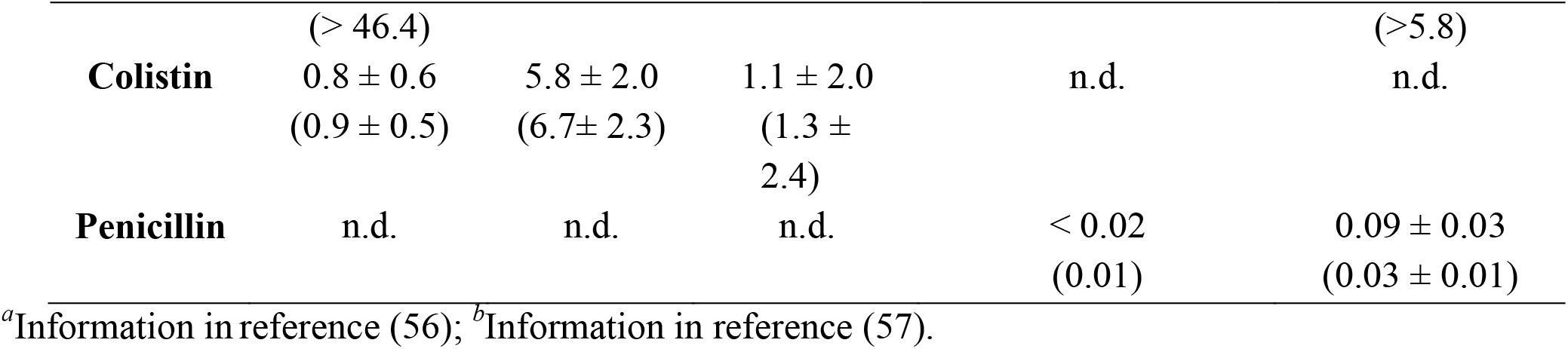
MIC values measured for the diruthenium compounds **1-22**, the ligand **23**, and colistin and penicillin used as controls. The reported values are the means of three biological replicates ± standard deviation. “n.d.” stands for “not determined”. The lowest MIC values for each strain are highlighted in bold. The values in brackets are the values expressed in μg/mL.

### 2.4. Determination of the MIC values on S. pneumoniae

In a second screening assay, the MIC values of diruthenium complexes **1-12** on *S. pneumoniae* D39 strain were determined, proving to be significantly lower compared to those measured on the three *E. coli* strains (Table 1). Compounds **5** and **6** were the most effective (MIC value of 1.3 μM for both compounds), slightly more active than compounds **1-3** (MIC values of 1.8-2.6 μM). The MIC values corresponding to compounds **7** and **8** were at the limit of the concentration used for the assay (0.625-10 μM) (Table 1) and being more than 5 times higher compared to the MIC value of compounds **5** and **6**. From Family 1 the dithiolato derivative **4** exhibited the lowest activity on *S. pneumoniae* D39. The MIC values of compounds **9-12** (Family 2) on *S. pneumoniae* D39 were higher than those found trithiolato compounds **1-3** and **5-7**.

### 2.5. Determination of the MIC values on S. aureus

In the third screening, MIC values of the 22 compounds on *S. aureus* 20 strain were also evaluated (Table 1). In Family 1, compound **6**, with a MIC value of 2.5 μM, was the most potent, followed by compounds **3**, **5** and **7** (MIC values ranging from 4.4 to 5 μM). Hydroxy compound **1** was the least efficient (MIC values of 17.5 μM), followed by the amino derivative **2**, and the dithiolato compound **4** (MIC value of 12.5 μM for both). Interestingly, compared to **1,** slight structural modifications in compounds **7** (^*i*^Pr group instead of ^*t*^Bu as substituents on two of the bridging thiols) lead to an improvement of activity against *S. aureus* 20 (MIC values of 17.5 and 5.0 μM for **1** and **7**, respectively). A similar effect was observed in the case of compound **8** compared to **2** for which the presence of BF_4_-as counterion instead of Cl^-^ was associated to a decrease of the MIC value on *S. aureus* 20 from 12.5 to 6.7 μM for **2** and **8**, respectively).

Interestingly, for compounds **1** and **2**, important differences of activity are observed on the two tested Gram-positive bacteria *S. aureus* 20 and *S. pneumoniae* D39.

In Family 2, conjugates **9** and **10**, with an hexanoic acid residue attached on one of the bridging thiols *via* ester or amide bonds, showed poor activity on *S. aureus* 20 (MIC values of 40 and 80 μM, respectively). Interestingly, the acetyl-protected glucose and galactose hybrids **11** and **12** exhibited lower MIC values (11 μM and 8.3 μM, respectively), but are up to four times less efficient in inhibiting the bacteria growth compared to **6**.

All the diruthenium compounds conjugated to BODIPY fluorophores (Family 3, compounds **13-16**) exhibited low activity against *S. aureus* 20 (MIC values over 80 μM).

Contrasting MIC values were obtained for compounds **17-22** (Family 4). From compounds **17-19** bearing *ortho*-substituted ligands, derivative **17** with a diphenyl-(2-mercaptophenyl)phosphonate bridging ligand, exhibited reduced efficacy on *S. aureus* 20 (MIC value of 40 μM), while the boron containing compound **18** and aldehyde compound **19** exhibited significantly lower MIC values (8.8 μM and 15 μM, respectively). Compounds **20-22**, bearing at least one benzo-fused lactam unit on the bridging thiols, exhibited low MIC values of 3.8, 2.5 and 3.1 mM, respectively.

The antibacterial activity of ligand **23** (7-mercapto-1,3,4,5-tetrahydro-2*H*-benzo[*b*]azepin-2-one) was also evaluated on *S. aureus,* using three times the concentration employed in the tests with compound **21**, but no bacteria growth inhibition was measured within the concentration range of the assay (7.5-240 μM).

The average MIC values of the 22 diruthenium compounds on the tested bacteria as a function of their respective molecular weight are shown in Figure S10, as well as the linear regression for each bacterial species. Interestingly, Figure S10 suggests a slight inverse correlation between MIC values and molecular weight for all bacterial species investigated in the present study. While this trend hardly exists for the *E. coli* phenotype P53.1R (only 5 compounds are considered), it appears evident for *S. pneumoniae* D39 and especially for *S. aureus* 20).

### 2.6. Evaluation of the bacteriostatic and bactericidal properties

The bacteriostatic and bactericidal properties of selected diruthenium compounds **1-3**, **5-8** and **11-12** were investigated on *S. aureus* 20 (Figure 5). Interestingly, the compounds exhibited a significant faster effect compared to penicillin, even when the penicillin concentration was increased to 8 times its MIC value (250 ng/L). The sharp decrease of the optical density at 600 nm (OD_600_) after the addition of all diruthenium compounds and the complete absence of cell growth within 24 h suggest a bactericidal rather than bacteriostatic effect.

**Figure 5.**
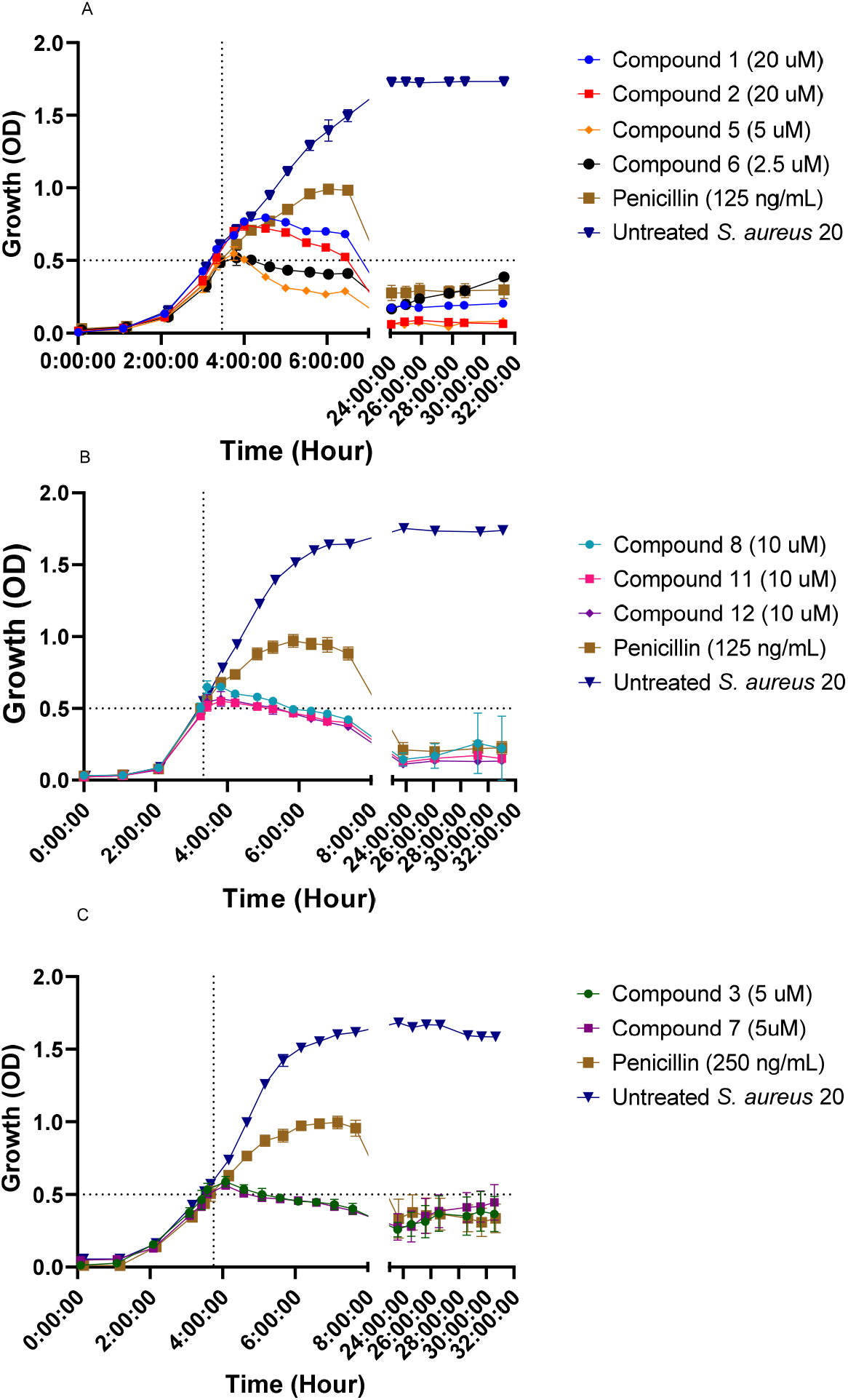
Growth curves of *S. aureus* 20 after addition of the diruthenium compounds at MIC concentration when OD_600_ = 0.5 is reached (dotted lines). **A**: *S. aureus* treated with compounds **1-2** and **5-6** at MIC concentrations upon reaching OD_600_ = 0.5. **B**: *S. aureus* treated with compounds **8**, **11** and **12** at MIC concentrations upon reaching OD_600_ = 0.5. **C**: *S. aureus* treated with compounds **3** and **7** at MIC concentrations upon reaching OD_600_ = 0.5. Treatment with penicillin at four times its MIC was used as a bactericidal control.

### 2.7. Fluorescence Microscopy

To determine whether the compounds preferentially accumulate inside the bacteria or on their membrane, conjugate **15** bearing a fluorescent BODIPY tag was used to investigate the localization of the diruthenium compounds by fluorescence microscopy. The hexanoic ester conjugate **9** was also tested in the fluorescence microscopy experiments as negative control (the fluorescence of **9** at 525/550 nm is shown in Figure 6, panels B). In parallel with compounds **9** and **15**, DAPI (4,6-diamidin-2-phenylindol) stain was used for the visualization of nuclear DNA.

**Figure 6.**
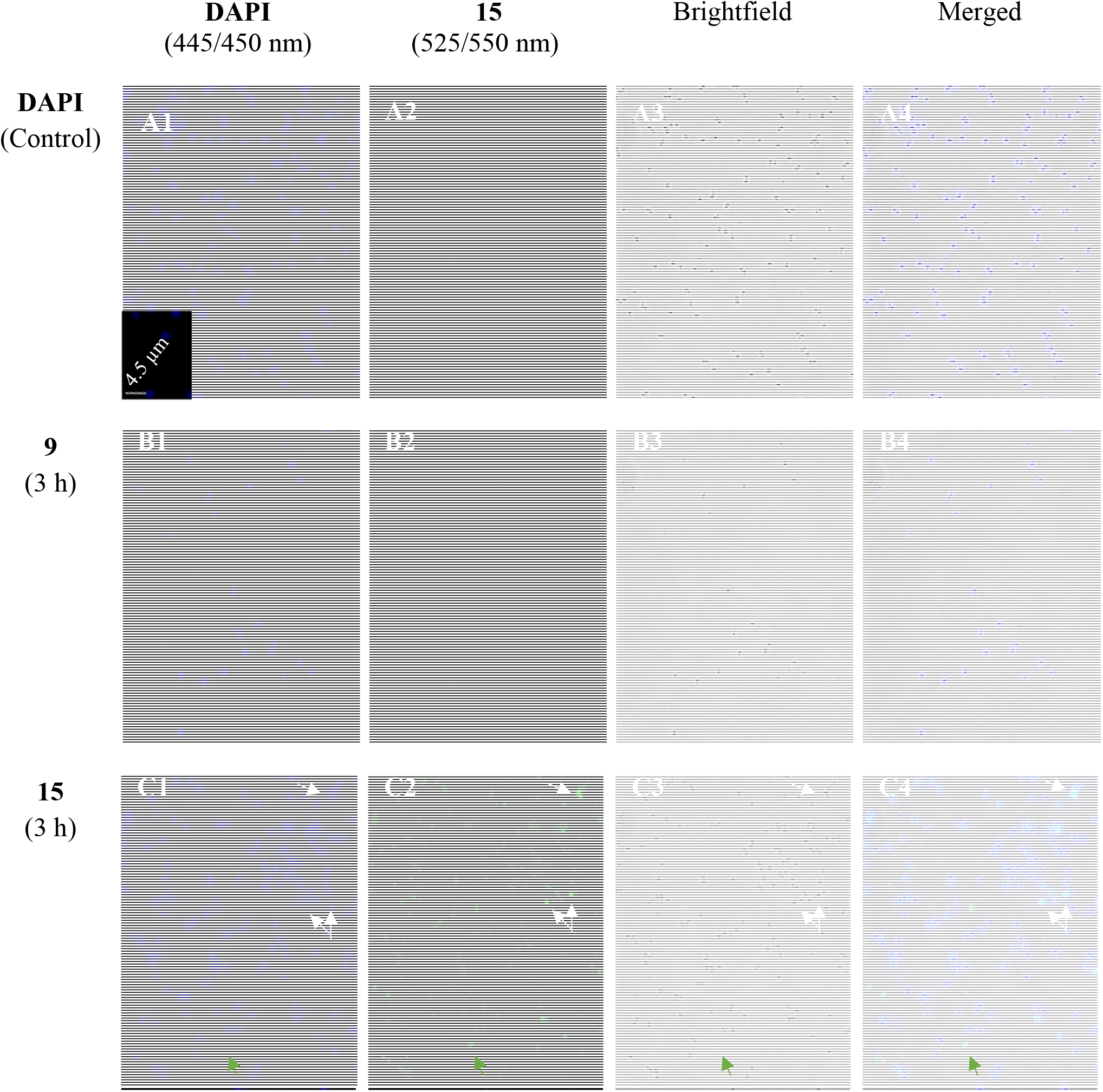
Fluorescence microscopy images of *S. aureus* 20 cells stained with DAPI after 3 h of incubation, with or without treatment with diruthenium conjugates **9** and **15**. A: Untreated control sample stained with DAPI. B: Sample treated with **9** for 3 h and stained with DAPI for visualization. C: Sample treated with **15** for 3 h and stained with DAPI for visualization. **15** is visible after excitation at 440/470 nm with emission at 525/550 nm. White arrows indicate cells with overlapped fluorescence of DAPI and BODIPY diruthenium conjugate **15**. Green arrows indicate emission in 525/550 nm (corresponding to conjugate **15**) which do not overlap with DAPI nor were cells visible under Brightfield.

In Figure 6 (panels C), an overlap of the blue fluorescent DAPI marker and the green fluorescence of BODIPY conjugate **15** inside the cells is clearly visible. Interestingly, emissions from **15** which do not overlap with those of DAPI are also observed (Figure 6, panels C). The results only validate the nuclear DNA as intracellular target of DAPI. The relative low fluorescence signal of compound **15** (52) does not allow to clearly distinguish the exact intracellular compound localization which might be the cytoplasm or the inner/outer membrane. Additional images are presented in Figures S12-S15 in *Supporting information.*

### 2.8. ICP-MS and cell counting

The cellular uptake of compounds **5**, **6**, **9**, **15** and **21** in the *S. aureus* 20 strain was quantified using ICP-MS measurements the results being summarized in Table 2. The bacteria were incubated with the selected ruthenium compounds at their respective MIC value for 1, 2, and 3 h. A clear increase of the ruthenium content of treated *S. aureus* 20 cells compared to the untreated control (mean value (6.6 ± 2.2)×10^-17^ μM/CFU(colony forming unit)) was measured, indicating cellular uptake of the diruthenium compounds (Table 2).

**Table 2.** Content of ruthenium in *S. aureus* 20 cells measured by ICP-MS. Values are the means of the 3 replicates ± standard deviation.

| Compound | Incubation time<br>(h) | Average compound concentration<br>per *S. aureus* cells ($\mu\text{M-CFU}$) |
| --- | --- | --- |
| **5** | 1 | $(7.7 \pm 2.0) \times 10^{-13}$ |
| | 2 | $(4.2 \pm 1.5) \times 10^{-13}$ |
| | 3 | $(4.1 \pm 1.3) \times 10^{-13}$ |
| **6** | 1 | $(4.0 \pm 1.2) \times 10^{-13}$ |
| | 2 | $(1.4 \pm 1.4) \times 10^{-12}$ |
| | 3 | $(1.4 \pm 1.1) \times 10^{-12}$ |
| **9** | 1 | $(2.8 \pm 1.9) \times 10^{-13}$ |
| | 2 | $(2.4 \pm 1.8) \times 10^{-13}$ |
| | 3 | $(2.6 \pm 1.3) \times 10^{-13}$ |
| **15** | 1 | $(2.2 \pm 1.7) \times 10^{-13}$ |
| | 2 | $(3.6 \pm 2.7) \times 10^{-13}$ |
| | 3 | $(3.4 \pm 2.6) \times 10^{-13}$ |
| **21** | 1 | $(6.6 \pm 0.6) \times 10^{-14}$ |
| | 2 | $(5.9 \pm 0.3) \times 10^{-14}$ |
| | 3 | $(6.1 \pm 1.2) \times 10^{-14}$ |

The amount of compound internalized by the cells appears to be independent of the treatment duration (Table 2). If overall within the treatment time frame of 3 h, no significant time-dependent modifications were observed for the same compound, important variations between the tested compounds were noticed. For example, the measured ruthenium concentration in *S. aureus* 20 cells after treatment with compound **6** (1.4×10^-12^ μM/CFU) was more than twenty times higher compared with the value for compound **21** (6×10^-14^ μM/CFU).

The results suggest a possible correlation between the amount of diruthenium compound internalized by the bacteria and the molecular weight, but no dependence on the respective MIC values (Figure S11). Compound **21**, bearing three 7-mercapto-1,3,4,5-tetrahydro-2*H*-benzo[*b*]azepin-2-one ligands as bridging thiols, is particularly interesting. **21** exhibits a low MIC value (2.5 μM, Table 1) but less efficient cellular uptake. These data suggest that **21** is highly toxic to the bacteria and this compound can be considered as a lead for further compound development through appropriate chemical modifications.

## 3. Discussion

In this work, the *in vitro* properties of a library of 22 trithiolato-bridged dinuclear ruthenium(II)-arene complexes as potential antibacterial agents against relevant bacterial species were investigated. The *in vitro* antiparasitic and anticancer activity of compounds **1-16** have been previously studied (40, 41, 44, 58). This type of diruthenium complexes are known to target in the mitochondria (40, 41, 44, 45, 59) an organelle sharing many features with bacterial cells (60).

Following the trend of most antibiotics (being more effective against Gram-positive than on Gram-negative pathogens (8, 12)), the results of this study revealed that the MIC values of the 22 tested compounds were higher on *E. coli* strains than on *S. pneumoniae* and *S. aureus*. Moreover, compared to colistin and penicillin tested as reference drugs, even the most effective compounds in this library were only moderately active, showing MIC values more than ten times higher.

For comparison, the MIC values of the ruthenium compounds *cis*-α-[Ru(phen)bb_12_]^2+^ and *cis-β-* [Ru(phen)bb_12_]^2+^ against the *S. aureus* MSSA ATCC 25,923 strain were 0.5 μM, and against the *E. coli* ATCC 25922 strain they were 8.3 and 16 μM, respectively (24). On the other hand, the MIC values of the ruthenium compound [Ru(bpy)_2_(methionine)]^2+^ against the *S. aureus* MSSA ATCC 25,923 strain and against the *E. coli* ATCC 11303 strain were of 73.3 and 586.4 μM, respectively (21).

The physico-chemical properties of the bridging thiol ligands are very important, their influence appearing particularly important in compounds **20-22** bearing benzo-fused lactams substituents, which exhibited low MIC values, and to a lesser extent, for carbohydrate conjugates **11-12**. The neutral dithiolato complex **4** had medium MIC values against *E. coli* and *S. pneumoniae*, following the trend observed with other dithiolato complexes against cancer cells and parasites (40, 47, 58). The lack of activity against cancer cells and parasites was attributed to the lability of the chloride ligands and the general lower stability of dithiolato compounds (61, 62). Yet, the MIC value of **4** against *S. aureus* was relatively low and comparable to that of the mixed trithiolato compounds **1** and **2**, compounds that were otherwise much more active than dithiolato complexes against cancer cells and protozoan parasites (40, 63). This difference is appealing and needs further investigations.

The cellular uptake of the BODIPY-diruthenium conjugate **15** by *S. aureus* was confirmed by fluorescence microscopy investigations. The images presented in Figures S12-S15 (*Supporting information*), indicate that compound **15** does not have a specific cellular localization. Figure S15 clearly shows that compound **15** does not accumulate in membranes or cell walls, but concentrates inside cells, forming aggregates that potentially lead to cell death.

In conclusion, the results show that the lower molecular weight diruthenium compounds tend to have lower MIC values against Gram-positive bacteria, but the presence of specific substituents, especially the 1,3,4,5-tetrahydro-2*H*-benzo[*b*]azepin-2-one unit, on the bridge thiols strongly influence the antibacterial activity. The 7-mercapto-1,3,4,5-tetrahydro-2*H*-benzo[*b*]azepin-2-one is an interesting ligand to be considered for the development of other potent antibacterial metal complexes. Finally, the localization and the cellular uptake of the compounds assessed using fluorescence microscopy and, respectively, ICP-MS suggest that the cellular walls, as well as the nucleic acid or protein synthesis are not the main targets of this type of compounds.

## 5. Materials and Methods

### 5.1. Materials

All new diruthenium complexes were synthesized, analyzed and characterized at the Department of Chemistry, Biochemistry and Pharmaceutical Sciences of University of Bern according to the methods provided in the *Supporting information*. Colistin, penicillin G potassium salt and gentamicin sulphate were purchased from Sigma-Aldrich and used as received.

### 5.2. Instrumentation

The OD_600_ of cell cultures and of the tubes used for the assays were measured with a Helios Epsilon spectrophotometer. in the In the MIC assays, the absorbance values for wells plates were obtained using a Varioskan Flash spectral scanning multimode plate reader using the SkanIt software. Cell lysis was realized using a Branson Ultrasonics Sonifier 250. Fluorescence microscopy observations were conducted using a ZEISS AXIO Imager M1, fluorescence microscope. ICP-MS values were determined with a NexION 2000B ICP Mass Spectrometer using Ir-193 as internal standard.

### 5.3. Antibacterial activity

#### 5.3.1. Cell culture

In this work were used three extended spectrum cephalosporin resistant *E. coli* strains (P53.1R, P54.1T and P54.2R) with varied colistin sensitivity, a penicillin sensitive strain of *S. pneumoniae* (D39) and a penicillin sensitive strain of *S. aureus* (20). The *E. coli* and *S. aureus* strains were inoculated in Luria-Bertani agar (LBA) plates and incubated overnight at 37°C. *S. pneumoniae* was inoculated in Columbia blood sheep agar (CSBA) plates and incubated overnight at 37°C and 5% CO_2_. Individual colonies of *E. coli* were then subcultured overnight at 37°C under constant agitation in Luria-Bertani (LB) medium for initial screening, and in Brain Heart Infusion (BHI) for the other assays. After the incubation period, the bacterial cultures were adjusted to a final concentration of 2×10^7^ colony-forming units (CFU)/mL. Individual colonies of *S. pneumoniae* and *S. aureus* were subcultured in BHI at 37°C until they reached the appropriate concentrations of 1×10^8^ and 2×10^8^ CFU/mL, respectively. The final concentration for inoculation in the assays were 5×10^5^, 5×10^6^ and 1×10^7^ CFU/mL, respectively.

#### 5.3.2. Determination of the minimum inhibitory concentration (MIC)

The assays were conducted in sterile Nunclon® 96 microwell plates and Nunc® 384 well polystyrene plates purchased from Thermo Fisher Scientific. DMSO (dimethyl sulfoxide) stock solutions of the diruthenium complexes (1-5 mM) were diluted in H_2_O to concentrations ranging from 2 μM to 200 μM according to compounds’ potency and dispensed into the plates with the previously adjusted bacterial suspension. The plates with *E. coli* or *S. aureus* were then incubated at 37°C for 18 h, and then the optical density (OD) at 450 nm was measured using a Varioskan Flash, spectral scanning multimode plate reader. The plates with *S. pneumoniae* were incubated at 37°C for 22 h with agitation and measurement every 30 min. The MIC was determined as the lowest compound concentration showing a complete inhibition of visible bacterial growth. For comparison, all assays included antibiotic controls: colistin, 8 μg/mL (*E. coli* P53.1R and P54.2R), 64 μg/mL (*E. coli* P54.1T) and penicillin 0.5 μg/mL (*S. pneumoniae* and *S. aureus*), as well as DMSO controls to account for the lethality of the organic solvent.

#### 5.3.3. Determination of the bacteriostatic/bactericidal effects

The assay was conducted in test tubes read with a Helios Epsilon spectrophotometer. DMSO stock solutions of the diruthenium complexes were diluted in H_2_O to concentrations 50 times their MIC values, before being added into *S. aureus* bacterial suspension in the test tube at 1:50 of the total volume, thus reaching MIC concentration. The tubes were incubated at 37°C in a water bath for 32 h. At the beginning of the assay, the OD (optical density) of each sample was measured every hour until an OD_600_ = 0.5 was reached. At this point, the previously prepared solution of diruthenium complexes and antibiotics were quickly added to their respective tubes and the negative growth controls before continuing incubation. Subsequently, the OD was measured for all samples every half hour until a total incubation time of 8 h was reached. Between 8 h and 24 h of incubation, the OD was not measured, and then was measured sporadically between 24 and 32 h of incubation to observe potential changes until the end of the assay.

#### 5.3.4. Cell counting

First, in a test tube, 5 mL BHI were added to 266.6 μL of a suspension of *S. aureus* in BHI and incubated at 37°C until an OD_600_ = 0.5 was measured. Subsequently, sequential dilutions were prepared, by adding 900 μL of a 0.85% NaCl aqueous solution to 100 μL from the suspension of *S. aureus* in BHI. This dilution procedure was repeated until a dilution of 10^-9^ was reached. 100 μL of the resulting 10^-3^ to 10^-9^ solutions were then put on CSBA plates and incubated at 37°C and 5% CO_2_ for 18 h. The CFU/mL of the original suspension was then estimated by counting the number of colonies in each plate and using a calibration curve. Each measurement was realized in triplicate.

### 5.4. Fluorescence Microscopy

Samples were prepared by incubating *S. aureus* bacterial suspensions for 3 h at 37°C in a water bath. Then 5 μM of compounds **9** and **15** were added and the suspensions were further incubated for 1, 2 and 3 h in 5 mL of BHI. Next, the samples were sedimented by centrifugation at 1788*g* for 12 minutes at 4 °C and washed twice with 0.85% aqueous NaCl. Then, the pellets were re-suspended in 3 mL 0.85% aqueous NaCl and cooled on ice before adding 2 μL of DAPI (DAPI Solution (1 mg/mL), Thermo Scientific™, 4’,6-diamidino-2-phenylindole, a fluorescent stain that binds strongly to adenine–thymine-rich DNA regions) stock solution. The samples were finally studied using a fluorescence microscope with filters for DAPI (excitation 365 nm, emission 445/50), GFP (green fluorescent compound, excitation 440/70, emission 525/5) and brightfield. To ensure observation of the cell internal, Z-stacks pictures were taken with an interval of 320 nm, using an objective 100x, 0.065 μm/pixel.

### 5.5. ICP-MS

The samples were prepared by incubating *S. aureus* bacterial suspensions (~4.3 x 10^8^ CFU/mL) for 3 h at 37°C in a water bath. Then 1 mM solution of compounds **5**, **6**, **9**, **15** and **21** in DMSO were added and the suspensions were incubated for 1, 2 and 3 h. A 150 μg/mL penicillin solution in distilled water was used to ensure cell death. The samples were then sedimented by centrifugation at 1788*g* for 12 minutes at 4 °C and washed with 0.85% aqueous NaCl twice. Afterward, the pellets were resuspended in 3 mL H_2_O and frozen at −20 °C overnight before undergoing heat shock and sonication for cell lysis. The resulting samples were then filtered using 0.22 μm Millipore syringe filters and analyzed by ICP-MS.

### 5.6. Determination of cellular uptake of the ruthenium compounds

Using the data obtained from the cell counting and ICP-MS experiments, and assuming comparable losses during filtration for both samples and controls, the intracellular ruthenium content was calculated using Equation 1:

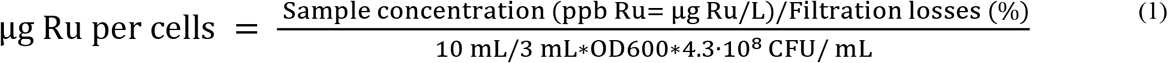

A calibration curve using ruthenium solutions of 0.02 ppb, 0.1 ppb, 1 ppb, 5 ppb and 50 ppb in 2% HNO3 was first established, and Iridium 193 was used as internal standard.

## Supplementary Materials

The following supporting information can be downloaded at:…, Chemistry, synthesis of compounds **17-22**, Characterization of the compounds, Figures **S1-S9**: NMR spectra; **Table S1:** Compilation of Silver-, Bismuth- and Ruthenium-based metal complexes investigated for their antibacterial properties; **Figure S10:** MIC values of the 22 diruthenium complexes as a function of their respective molecular weight. **Figure S11**: Cellular uptake of compounds **5, 6, 9, 15** and **21** as a function of the MIC values against S. aureus 20 and Cellular uptake of compounds **5, 6, 9, 15** and **21** as a function of the molecular weight. **Figure S12**: Z-cut photography under DAPI filter (445/450 nm) of *S. aureus* 20 cells treated with compound **15** for 3 h. **Figure S13:** Z-cut photography under GFP filter (525/550 nm) of *S. aureus* 20 cells treated with compound **15** for 3 h; **Figure S14:** Z-cut photography under Brightfield of *S. aureus* 20 cells treated with compound **15** for 3 h; **Figure S15**: Merged Z-cut photography with DAPI filter (445/450 nm), GFP filter (525/550 nm) and Brightfield of *S. aureus* 20 cells treated with compound **15** for 3 h.

## Data Availability Statement

All the data are presented in this study and in the corresponding *Supporting information.*

## References

1. Origins and Evolution of Antibiotic Resistance | Microbiology and Molecular Biology Reviews.

2. Frieri M, Kumar K, Boutin A. 2017.Antibiotic resistance. Journal of Infection and Public Health 10:369–378.

3. Jani K, Srivastava V, Sharma P, Vir A, Sharma A. 2021. Easy Access to Antibiotics; Spread of Antimicrobial Resistance and Implementation of One Health Approach in India. J Epidemiol Glob Health 11:444–452.

4. Terreni M, Taccani M, Pregnolato M. 2021. New Antibiotics for Multidrug-Resistant Bacterial Strains: Latest Research Developments and Future Perspectives. Molecules 26:2671.

5. Murray CJ, Ikuta KS, Sharara F, Swetschinski L, Aguilar GR, Gray A, Han C, Bisignano C, Rao P, Wool E, Johnson SC, Browne AJ, Chipeta MG, Fell F, Hackett S, Haines-Woodhouse G, Hamadani BHK, Kumaran EAP, McManigal B, Agarwal R, Akech S, Albertson S, Amuasi J, Andrews J, Aravkin A, Ashley E, Bailey F, Baker S, Basnyat B, Bekker A, Bender R, Bethou A, Bielicki J, Boonkasidecha S, Bukosia J, Carvalheiro C, Castañeda-Orjuela C, Chansamouth V, Chaurasia S, Chiurchiù S, Chowdhury F, Cook AJ, Cooper B, Cressey TR, Criollo-Mora E, Cunningham M, Darboe S, Day NPJ, Luca MD, Dokova K, Dramowski A, Dunachie SJ, Eckmanns T, Eibach D, Emami A, Feasey N, Fisher-Pearson N, Forrest K, Garrett D, Gastmeier P, Giref AZ, Greer RC, Gupta V, Haller S, Haselbeck A, Hay SI, Holm M, Hopkins S, Iregbu KC, Jacobs J, Jarovsky D, Javanmardi F, Khorana M, Kissoon N, Kobeissi E, Kostyanev T, Krapp F, Krumkamp R, Kumar A, Kyu HH, Lim C, Limmathurotsakul D, Loftus MJ, Lunn M, Ma J, Mturi N, Munera-Huertas T, Musicha P, Mussi-Pinhata MM, Nakamura T, Nanavati R, Nangia S, Newton P, Ngoun C, Novotney A, Nwakanma D, Obiero CW, Olivas-Martinez A, Olliaro P, Ooko E, Ortiz-Brizuela E, Peleg AY, Perrone C, Plakkal N, Ponce-de-Leon A, Raad M, Ramdin T, Riddell A, Roberts T, Robotham JV, Roca A, Rudd KE, Russell N, Schnall J, Scott JAG, Shivamallappa M, Sifuentes-Osornio J, Steenkeste N, Stewardson AJ, Stoeva T, Tasak N, Thaiprakong A, Thwaites G, Turner C, Turner P, Doorn HR van, Velaphi S, Vongpradith A, Vu H, Walsh T, Waner S, Wangrangsimakul T, Wozniak T, Zheng P, Sartorius B, Lopez AD, Stergachis A, Moore C, Dolecek C, Naghavi M. 2022. Global burden of bacterial antimicrobial resistance in 2019: a systematic analysis. The Lancet 399:629–655.

6. Regea G. 2018. Pharmacology & Clinical Research Review on Antibiotics Resistance and its Economic Impacts https://doi.org/10.19080/JPCR.2018.05.555675.

7. Tacconelli E, Carrara E, Savoldi A, Harbarth S, Mendelson M, Monnet DL, Pulcini C, Kahlmeter G, Kluytmans J, Carmeli Y, Ouellette M, Outterson K, Patel J, Cavaleri M, Cox EM, Houchens CR, Grayson ML, Hansen P, Singh N, Theuretzbacher U, Magrini N, WHO Pathogens Priority List Working Group. 2018. Discovery, research, and development of new antibiotics: the WHO priority list of antibiotic-resistant bacteria and tuberculosis. Lancet Infect Dis 18:318–327.

8. Breijyeh Z, Jubeh B, Karaman R. 2020. Resistance of Gram-Negative Bacteria to Current Antibacterial Agents and Approaches to Resolve It. Molecules 25:E1340.

9. Mitcheltree MJ, Pisipati A, Syroegin EA, Silvestre KJ, Klepacki D, Mason JD, Terwilliger DW, Testolin G, Pote AR, Wu KJY, Ladley RP, Chatman K, Mankin AS, Polikanov YS, Myers AG. 2021. A synthetic antibiotic class overcoming bacterial multidrug resistance. 7885. Nature 599:507–512.

10. Frei A, Zuegg J, Elliott AG, Baker M, Braese S, Brown C, Chen F, Dowson CG, Dujardin G, Jung N, King AP, Mansour AM, Massi M, Moat J, Mohamed HA, Renfrew AK, Rutledge PJ, Sadler PJ, Todd MH, Willans CE, Wilson JJ, Cooper MA, Blaskovich MAT. 2020. Metal complexes as a promising source for new antibiotics. Chem Sci 11:2627–2639.

11. Gasser G. 2015. Metal Complexes and Medicine: A Successful Combination. Chimia (Aarau) 69:442–446.

12. Munteanu A-C, Uivarosi V. 2021. Ruthenium Complexes in the Fight against Pathogenic Microorganisms. An Extensive Review. 6. Pharmaceutics 13:874.

13. Sharkey MA, O’Gara JP, Gordon SV, Hackenberg F, Healy C, Paradisi F, Patil S, Schaible B, Tacke M. 2012. Investigations into the Antibacterial Activity of the Silver-Based Antibiotic Drug Candidate SBC3. Antibiotics (Basel) 1:25–28.

14. Haziz UFM, Haque RA, Amirul AA, Razali MR. 2021. Synthesis, Structural Analysis and Antibacterial Studies of Bis-and Open Chain Tetra-N-Heterocyclic Carbene Dinuclear Silver(I) Complexes. Journal of Molecular Structure 1236:130301.

15. López-Cardoso M, Tlahuext H, Pérez-Salgado M, Vargas-Pineda DG, Román-Bravo PP, Cotero-Villegas AM, Acevedo-Quiroz M, Razo-Hernández RS, Alvarez-Fitz P, Mendoza-Catalán MA, Jancik V, Cea-Olivares R. 2020. Synthesis, crystal structure, antibacterial, antiproliferative and QSAR studies of new bismuth(III) complexes of pyrrolidineditiocarbamate of dithia-bismolane and bismane, oxodithia-and trithia-bismocane. Journal of Molecular Structure 1217:128456.

16. Marzano IM, Tomco D, Staples RJ, Lizarazo-Jaimes EH, Gomes DA, Bucciarelli-Rodriguez M, Guerra W, de Souza ÍP, Verani CN, Pereira Maia EC. 2021. Dual anticancer and antibacterial activities of bismuth compounds based on asymmetric [NN’O] ligands. Journal of Inorganic Biochemistry 222:111522.

17. Kollipara MR, Shadap L, Banothu V, Agarwal N, Poluri KM, Kaminsky W. 2020. Fluorenone Schiff base derivative complexes of ruthenium, rhodium and iridium exhibiting efficient antibacterial activity and DNA-binding affinity. Journal of Organometallic Chemistry 915:121246.

18. Nolan VC, Rafols L, Harrison J, Soldevila-Barreda JJ, Crosatti M, Garton NJ, Wegrzyn M, Timms DL, Seaton CC, Sendron H, Azmanova M, Barry NPE, Pitto-Barry A, Cox JAG. 2022. Indole-containing arene-ruthenium complexes with broad spectrum activity against antibiotic-resistant bacteria. Current Research in Microbial Sciences 3:100099.

19. Bolhuis A, Hand L, Marshall JE, Richards AD, Rodger A, Aldrich-Wright J. 2011. Antimicrobial activity of ruthenium-based intercalators. European Journal of Pharmaceutical Sciences 42:313–317.

20. Yang X-Y, Zhang L, Liu J, Li N, Yu G, Cao K, Han J, Zeng G, Pan Y, Sun X, He Q-Y. 2015. Proteomic analysis on the antibacterial activity of a Ru(II) complex against Streptococcus pneumoniae. J Proteomics 115:107–116.

21. de Sousa AP, Gondim ACS, Sousa EHS, de Vasconcelos MA, Teixeira EH, Bezerra BP, Ayala AP, Martins PHR, Lopes LG de F, Holanda AKM. 2020. An unusual bidentate methionine ruthenium(II) complex: photo-uncaging and antimicrobial activity. J Biol Inorg Chem 25:419–428.

22. Srivastava P, Shukla M, Kaul G, Chopra S, Patra AK. 2019. Rationally designed curcumin based ruthenium(II) antimicrobials effective against drug-resistant Staphylococcus aureus. Dalton Trans 48:11822–11828.

23. Li F, Feterl M, Mulyana Y, Warner JM, Collins JG, Keene FR. 2012. In vitro susceptibility and cellular uptake for a new class of antimicrobial agents: dinuclear ruthenium(II) complexes. J Antimicrob Chemother 67:2686–2695.

24. Gorle AK, Feterl M, Warner JM, Primrose S, Constantinoiu CC, Keene FR, Collins JG. 2015. Mononuclear Polypyridylruthenium(II) Complexes with High Membrane Permeability in Gram-Negative Bacteria-in particular Pseudomonas aeruginosa. Chemistry 21:10472–10481.

25. Smitten KL, Thick EJ, Southam HM, Serna JB de la, Foster SJ, Thomas JA. 2020. Mononuclear ruthenium(II) theranostic complexes that function as broad-spectrum antimicrobials in therapeutically resistant pathogens through interaction with DNA. Chem Sci 11:8828–8838.

26. Kumar SV, Scottwell SØ, Waugh E, McAdam CJ, Hanton LR, Brooks HJL, Crowley JD. 2016. Antimicrobial Properties of Tris(homoleptic) Ruthenium(II) 2-Pyridyl-1,2,3-triazole “Click” Complexes against Pathogenic Bacteria, Including Methicillin-Resistant Staphylococcus aureus (MRSA). Inorg Chem 55:9767–9777.

27. van Hilst QVC, Vasdev RAS, Preston D, Findlay JA, Scottwell SØ, Giles GI, Brooks HJL, Crowley JD. 2019. Synthesis, Characterisation and Antimicrobial Studies of some 2,6-bis(1,2,3-Triazol-4-yl)Pyridine Ruthenium(II) “Click” Complexes. Asian Journal of Organic Chemistry 8:496–505.

28. Li F, Mulyana Y, Feterl M, Warner JM, Collins JG, Keene FR. 2011. The antimicrobial activity of inert oligonuclear polypyridylruthenium(II) complexes against pathogenic bacteria, including MRSA. Dalton Trans 40:5032–5038.

29. Smitten KL, Southam HM, de la Serna JB, Gill MR, Jarman PJ, Smythe CGW, Poole RK, Thomas JA. 2019. Using Nanoscopy To Probe the Biological Activity of Antimicrobial Leads That Display Potent Activity against Pathogenic, Multidrug Resistant, Gram-Negative Bacteria. ACS Nano 13:5133–5146.

30. Smitten KL, Fairbanks SD, Robertson CC, Serna JB de la, Foster SJ, Thomas JA. 2019. Ruthenium based antimicrobial theranostics – using nanoscopy to identify therapeutic targets and resistance mechanisms in Staphylococcus aureus. Chem Sci 11:70–79.

31. Pandrala M, Li F, Feterl M, Mulyana Y, Warner JM, Wallace L, Keene FR, Collins JG. 2013. Chlorido-containing ruthenium(II) and iridium(III) complexes as antimicrobial agents. Dalton Trans 42:4686–4694.

32. Sun B, Sundaraneedi MK, Southam HM, Poole RK, Musgrave IF, Keene FR, Collins JG. 2019. Synthesis and biological properties of tetranuclear ruthenium complexes containing the bis[4(4’-methyl-2,2’-bipyridyl)]-1,7-heptane ligand. Dalton Trans 48:14505–14515.

33. Mi H, Wang D, Xue Y, Zhang Z, Niu J, Hong Y, Drlica K, Zhao X. 2016. Dimethyl Sulfoxide Protects Escherichia coli from Rapid Antimicrobial-Mediated Killing. Antimicrob Agents Chemother 60:5054–5058.

34. Smith H, Mann BE, Motterlini R, Poole RK. 2011. The carbon monoxide-releasing molecule, corm-3 (ru(co)3cl(glycinate)), targets respiration and oxidases in campylobacter jejuni, generating hydrogen peroxide. IUBMB Life 63:363–371.

35. Wilson JL, Jesse HE, Hughes B, Lund V, Naylor K, Davidge KS, Cook GM, Mann BE, Poole RK. 2013. Ru(CO)3Cl(Glycinate) (CORM-3): A Carbon Monoxide–Releasing Molecule with Broad-Spectrum Antimicrobial and Photosensitive Activities Against Respiration and Cation Transport in Escherichia coli. Antioxidants & Redox Signaling 19:497–509.

36. Wareham LK, Poole RK, Tinajero-Trejo M. 2015. CO-releasing Metal Carbonyl Compounds as Antimicrobial Agents in the Post-antibiotic Era. J Biol Chem 290:18999–19007.

37. Furrer J, Süss-Fink G. 2016. Thiolato-bridged dinuclear arene ruthenium complexes and their potential as anticancer drugs. Coordination Chemistry Reviews 309:36–50.

38. Giannini F, Süss-Fink G, Furrer J. 2011. Efficient oxidation of cysteine and glutathione catalyzed by a dinuclear areneruthenium trithiolato anticancer complex. Inorganic Chemistry 50:10552–10554.

39. Tomšík P, Muthná D, Řezáčová M, Mičuda S, Ćmielová J, Hroch M, Endlicher R, Červinková Z, Rudolf E, Hann S, Stíbal D, Therrien B, Süss-Fink G. 2015. [(p-MeC6H4Pri)2Ru2(SC6H4-p-But)3]Cl (diruthenium-1), a dinuclear arene ruthenium compound with very high anticancer activity: An in vitro and in vivo study. Journal of Organometallic Chemistry 782:42–51.

40. Basto AP, Müller J, Rubbiani R, Stibal D, Giannini F, Süss-Fink G, Balmer V, Hemphill A, Gasser G, Furrer J. 2017. Characterization of the activities of dinuclear thiolato-bridged arene ruthenium complexes against Toxoplasma gondii. Antimicrobial Agents and Chemotherapy 61.

41. Basto AP, Anghel N, Rubbiani R, Müller J, Stibal D, Giannini F, Süss-Fink G, Balmer V, Gasser G, Furrer J, Hemphill A. 2019. Targeting of the mitochondrion by dinuclear thiolato-bridged arene ruthenium complexes in cancer cells and in the apicomplexan parasite Neospora caninum. Metallomics 11:462–474.

42. Desiatkina O, Păunescu E, Mösching M, Anghel N, Boubaker G, Amdouni Y, Hemphill A, Furrer J. 2020. Coumarin-Tagged Dinuclear Trithiolato-Bridged Ruthenium(II)·Arene Complexes: Photophysical Properties and Antiparasitic Activity. ChemBioChem 21:2818–2835.

43. Desiatkina O, Johns SK, Anghel N, Boubaker G, Hemphill A, Furrer J, Păunescu E. 2021. Synthesis and Antiparasitic Activity of New Conjugates—Organic Drugs Tethered to Trithiolato-Bridged Dinuclear Ruthenium(II)–Arene Complexes. 8. Inorganics 9:59.

44. Paunescu E, Boubaker G, Desiatkina O, Anghel N, Amdouni Y, Hemphill A, Furrer J. 2021. The quest of the best - A SAR study of trithiolato-bridged dinuclear Ruthenium(II)-Arene compounds presenting antiparasitic properties. Eur J Med Chem 222:113610.

45. Jelk J, Balmer V, Stibal D, Giannini F, Süss-Fink G, Bütikofer P, Furrer J, Hemphill A. 2019. Anti-parasitic dinuclear thiolato-bridged arene ruthenium complexes alter the mitochondrial ultrastructure and membrane potential in Trypanosoma brucei bloodstream forms. Experimental Parasitology 205.

46. Giannini F, Paul LEH, Furrer J, Therrien B, Süss-Fink G. 2013. Highly cytotoxic diruthenium trithiolato complexes of the type [(η6-p-MeC6H4Pri) 2Ru2(μ2-SR)3]+: Synthesis, characterization, molecular structure and in vitro anticancer activity. New Journal of Chemistry 37:3503–3511.

47. Ibao A-F, Gras M, Therrien B, Süss-Fink G, Zava O, Dyson PJ. 2012. Thiolato-Bridged Arene–Ruthenium Complexes: Synthesis, Molecular Structure, Reactivity, and Anticancer Activity of the Dinuclear Complexes [(arene)2Ru2(SR)2Cl2]. European Journal of Inorganic Chemistry 2012:1531–1535.

48. Desiatkina O, Anghel N, Boubaker G, Amdouni Y, Hemphill A, Furrer J, Păunescu E. 2023. Trithiolato-bridged dinuclear ruthenium(II)-arene conjugates tethered with lipophilic units: Synthesis and Toxoplasma gondii antiparasitic activity. Journal of Organometallic Chemistry 986:122624.

49. Holzer I, Desiatkina O, Anghel N, Johns SK, Boubaker G, Hemphill A, Furrer J, Paunescu E. 2022. Synthesis and Antiparasitic Activity of New Trithiolato-Bridged Dinuclear Arene Ruthenium-Glycoconjugates submitted to Molecules.

50. Desiatkina O, Anghel N, Boubaker G, Amdouni Y, Hemphill A, Furrer J, Paunescu E. 2022. Trithiolato-Bridged Dinuclear Ruthenium(II)-Arene Conjugates Tethered with Lipophilic Units: Synthesis and Antiparasitic Activity manuscript in preparation.

51. Bertrand B, Passador K, Goze C, Denat F, Bodio E, Salmain M. 2018. Metal-based BODIPY derivatives as multimodal tools for life sciences. Coordination Chemistry Reviews 358:108–124.

52. Desiatkina O, Boubaker G, Anghel N, Amdouni Y, Hemphill A, Furrer J, Păunescu E. Synthesis, Photophysical Properties and Biological Evaluation of New Conjugates BODIPY: Dinuclear Trithiolato-Bridged Ruthenium(II)-Arene Complexes. ChemBioChem n/a:e202200536.

53. Figuly GD, Loop CK, Martin JC. 1989. Directed ortho-lithiation of lithium thiophenolate. New methodology for the preparation of ortho-substituted thiophenols and related compounds. J Am Chem Soc 111:654–658.

54. Wang L-Z, Li X-Q, An Y-S. 2015. 1,5-Benzodiazepine derivatives as potential antimicrobial agents: design, synthesis, biological evaluation, and structure–activity relationships. Org Biomol Chem 13:5497–5509.

55. An Y, Hao Z, Zhang X, Wang L. 2016. Efficient Synthesis and Biological Evaluation of a Novel Series of 1,5-Benzodiazepine Derivatives as Potential Antimicrobial Agents. Chemical Biology & Drug Design 88:110–121.

56. Sudatip D, Tiengrim S, Chasiri K, Kritiyakan A, Phanprasit W, Morand S, Thamlikitkul V. 2022. *One Health* Surveillance of Antimicrobial Resistance Phenotypes in Selected Communities in Thailand. Antibiotics (Basel) 11:556.

57. Iannelli F, Pearce BJ, Pozzi G. 1999. The type 2 capsule locus of Streptococcus pneumoniae. J Bacteriol 181:2652–2654.

58. Furrer J, Süss-Fink G. 2016. Thiolato-bridged dinuclear arene ruthenium complexes and their potential as anticancer drugs. Coordination Chemistry Reviews 309:36–50.

59. Anghel N, Müller J, Serricchio M, Jelk J, Bütikofer P, Boubaker G, Imhof D, Ramseier J, Desiatkina O, Păunescu E, Braga-Lagache S, Heller M, Furrer J, Hemphill A. 2021. Cellular and molecular targets of nucleotide-tagged trithiolato-bridged arene ruthenium complexes in the protozoan parasites toxoplasma gondii and trypanosoma brucei. International Journal of Molecular Sciences 22.

60. Boguszewska K, Szewczuk M, Kaźmierczak-Barańska J, Karwowski BT. 2020. The Similarities between Human Mitochondria and Bacteria in the Context of Structure, Genome, and Base Excision Repair System. Molecules 25:2857.

61. Stíbal D, Geiser L, Süss-Fink G, Furrer J. 2016. Hydrolytic behaviour of mono-and dithiolato-bridged dinuclear arene ruthenium complexes and their interactions with biological ligands. RSC Adv 6:38332–38341.

62. Primasová H, Ninova S, Capitani M de, Daepp J, Aschauer U, Furrer J. 2020. Dinuclear thiolato-bridged arene ruthenium complexes: from reaction conditions and mechanism to synthesis of new complexes. RSC Adv 10:40106–40116.

63. Giannini F, Furrer J, Ibao A-F, Süss-Fink G, Therrien B, Zava O, Baquie M, Dyson PJ, Štĕpnička P. 2012. Highly cytotoxic trithiophenolatodiruthenium complexes of the type [(η6-p-MeC6H4Pr i) 2Ru2(SC6H4-p-X)3] +: Synthesis, molecular structure, electrochemistry, cytotoxicity, and glutathione oxidation potential. Journal of Biological Inorganic Chemistry 17:951–960.

